# A merger between compatible but divergent genomes supports allopolyploidization in the Brassicaceae family

**DOI:** 10.1101/2021.09.28.462158

**Authors:** Hosub Shin, Jeong Eun Park, Hye Rang Park, Woo Lee Choi, Seung Hwa Yu, Wonjun Koh, Seungill Kim, Hye Yeon Soh, Nomar Espinosa Waminal, Hadassah Roa Belandres, Joo Young Lim, Gibum Yi, Jong Hwa Ahn, June-Sik Kim, Yong-Min Kim, Namjin Koo, Kyunghee Kim, Sampath Perumal, Taegu Kang, Junghyo Kim, Hosung Jang, Dong Hyun Kang, Ye Seul Kim, Hyeon-Min Jeong, Junwoo Yang, Somin Song, Suhyoung Park, Jin A Kim, Yong Pyo Lim, Beom-Seok Park, Tzung-Fu Hsieh, Tae-Jin Yang, Doil Choi, Hyun Hee Kim, Soo-Seong Lee, Jin Hoe Huh

## Abstract

Hybridization and polyploidization are pivotal to plant evolution. Genetic crosses between distantly related species rarely occur in nature mainly due to reproductive barriers but how such hurdles can be overcome is largely unknown. x*Brassicoraphanus* is a fertile intergeneric allopolyploid synthesized between *Brassica rapa* and *Raphanus sativus* in the Brassicaceae family. Genomes of *B. rapa* and *R. sativus* are diverged enough to suppress synapsis formation between non-homologous progenitor chromosomes during meiosis, and we found that both genomes reside in the single nucleus of x*Brassicoraphanus* without genome loss or rearrangement. Expressions of syntenic orthologs identified in *B. rapa* and *R. sativus* were adjusted to a hybrid nuclear environment of x*Brassicoraphanus*, which necessitates reconfiguration of transcription network by rewiring *cis*-*trans* interactions. *B. rapa* coding sequences have a higher level of gene-body methylation than *R. sativus*, and such methylation asymmetry is maintained in x*Brassicoraphanus*. *B. rapa*-originated transposable elements were transcriptionally silenced in x*Brassicoraphanus*, rendered by gain of CHG methylation *in trans* via small RNAs derived from the same sequences of *R. sativus* subgenome. Our work proposes that not only transcription compatibility but also a certain extent of genome divergence supports hybrid genome stabilization, which may explain great diversification and expansion of angiosperms during evolution.

## Introduction

Genome hybridization and polyploidization have served as major driving forces in plant evolution (Wendel, 2000; Soltis and Soltis, 2009, 2016; Van de Peer et al., 2017; Cheng et al., 2018). However, strong hybridization barriers exist in nature to prevent a gene flow between different species in plants and animals (Abbott et al., 2013). Several mechanisms have been proposed to explain the postzygotic barriers resulting from genome incompatibility between distantly related species (Lafon-Placette and Kohler, 2015; Dion-Cote and Barbash, 2017). Among them, a ‘genome shock’ is proposed as one of the critical causes of genome destabilization upon hybridization, restructuring the hybrid genome through changes of chromosomal organization or mobilization of transposable elements (TEs) (McClintock, 1984). Another is a ‘transcriptome shock’ that incurs extensive changes of parental gene expression patterns in the hybrid (Hegarty et al., 2006; Buggs et al., 2011).

Despite such negative consequences of hybridization between distantly related species, novel species can be naturally or artificially produced in a rare occasion while overcoming the hybridization barrier, the mechanism of which is largely unknown. The Brassicaceae family contains a variety of agronomically important crop species such as broccoli, cabbage, cauliflower, oilseed rape, radish and turnip, in addition to a model plant *Arabidopsis*. The genus *Brassica* is well known for hybridization between different species within the same genus (interspecific hybridization). For instance, three diploid species *Brassica rapa* (*Br*; AA), *B. nigra* (BB) and *B. oleracea* (*Bo*; CC) can hybridize each other generating allotetraploid species *B. napus* (AACC), *B. juncea* (AABB) and *B. carinata* (BBCC), as epitomized by the model of ‘Triangle of U’ (U, 1935).

Hybridization between species in the Brassicaceae family is not restricted to interspecific hybridization. Since 1826 by Sageret (Oost, 1984), intergeneric hybrids between *Brassica* and *Raphanus* have been sporadically reported (Karpechenko, 1928; Mcnaughton, 1973; Dolstra, 1982) but failed to survive. Recently developed x*Brassicoraphanus* (x*B*) (AARR; 2n = 4x = 38) is an intergeneric allotetraploid between *Br* (AA; 2n = 2x = 20) and *Raphanus sativus* (*Rs*) (RR; 2n = 2x = 18) (Lee et al., 2011). Unlike most newly synthesized interspecific/intergeneric hybrids, x*B* is self-fertile and genetically stable displaying phenotypic uniformity in successive generations (Supplemental Figure S1). Genetic and phenotypic stability of x*B* is very exceptional considering that many allopolyploids often display a high degree of genome instability and sterility issues, indicating that the hybridization barrier was overcome immediately after the two genomes have merged.

We hypothesized that allopolyploidization events have somewhat ameliorated deleterious shock phenomena such as genome and transcriptome shocks, and thereby overcome an intrinsic hybridization barrier between distantly related species. We here report genome structure, chromosome behaviors, and transcriptome/epigenome profiles of x*B*. We observed inhibition of meiotic non-homologous interactions, adjustment of homoeologous gene expressions and gain of DNA methylation. All these likely contribute to genome stability and transcription network compatibility in x*B*. This study further proposes the possible mechanisms by which two divergent genomes can successfully merge into a novel species during evolution of angiosperms.

## Results

### Genomic features of x*Brassicoraphanus*

x*B* is a fertile and genetically stable intergeneric allotetraploid synthesized from a cross between *Br* and *Rs*. The x*B* genome was *de novo* assembled using 195.0 Gb of Illumina shotgun reads (Figure 1A, Table 1 and Supplemental Tables S1 and S2). Flow cytometry analysis estimated the size of x*B* genome as 998.3 Mb, close to the sum of *Br* (485 Mb) and *Rs* (510 Mb) genomes (Wang et al., 2011; Jeong et al., 2016) (Supplemental Figure S2). We assembled 692.8 Mb sequence covering ∼70% of the x*B* genome, which contains 87,861 annotated genes and 39.19% (255.8 Mb) of repeat regions with long terminal repeats (LTRs) being predominant (Supplemental Table S3). The assembled chloroplast genome of x*B* (153,482 bp) was 99.9% identical to that of *Br* indicating its maternal origin (Supplemental Figure S3 and Supplemental Table S4). In x*B* genome (692.8 Mb), 335.5 Mb and 343.5 Mb of scaffolds were assigned to *Br* and *Rs* genomes (referred to as A_Br_ and R_Rs_ hereafter), respectively (Wang et al., 2011; Jeong et al., 2016), comprising two subgenomes of x*B* (referred to as A_xB_ and R_xB_ hereafter) (Table 1 and Supplemental Figure S4). Differentially expressed genes (DEGs) whose expressions are up- or down-regulated relative to the progenitors emerge evenly throughout the x*B* genome (Figure 1A). DNA methylation is predominant in repeat-enriched regions at all CG, CHG and CHH (H = A, T or C) contexts (Figure 1A and Supplemental Figure S5). Differentially methylated regions (DMRs) refer to the regions where DNA methylation levels in x*B* are significantly different (absolute difference > 0.3 for CG, > 0.15 for CHG and > 0.1 for CHH) from those of *Br* and *Rs*, and about 60.2% of hyper-DMRs are confined to repeat regions (Supplemental Figure S5 and Supplemental Data Set S1). Approximately 75.8% of H3K9me2 repressive histone marks are also enriched in repeat regions (Figure 1A, Supplemental Figure S6 and Supplemental Data Set S2). Small RNAs (18-30 nt) are distributed throughout the entire x*B* genome and significantly associated with DNA methylation (Figure 1A). Cytological observation revealed a total of 19 chromosome pairs present in x*B* without aneuploidy and/or chromosome rearrangements (Figure 1B). Previous studies reported that many synthetic allopolyploid plants such as rapeseed, tobacco and wheat went through massive chromosome reconstruction leading to transgressive gain or loss of chromosomes and/or aneuploidy over generations (Xiong et al., 2011; Zhang et al., 2013; Chen et al., 2018; Sosnowska et al., 2020). However, our findings indicate that x*B* retains both A_Br_ and R_Rs_ genomes in the single nucleus without structural aberrations, but at the same time, experiences substantial changes in transcriptome and epigenome profiles after hybridization.

**Figure 1.**
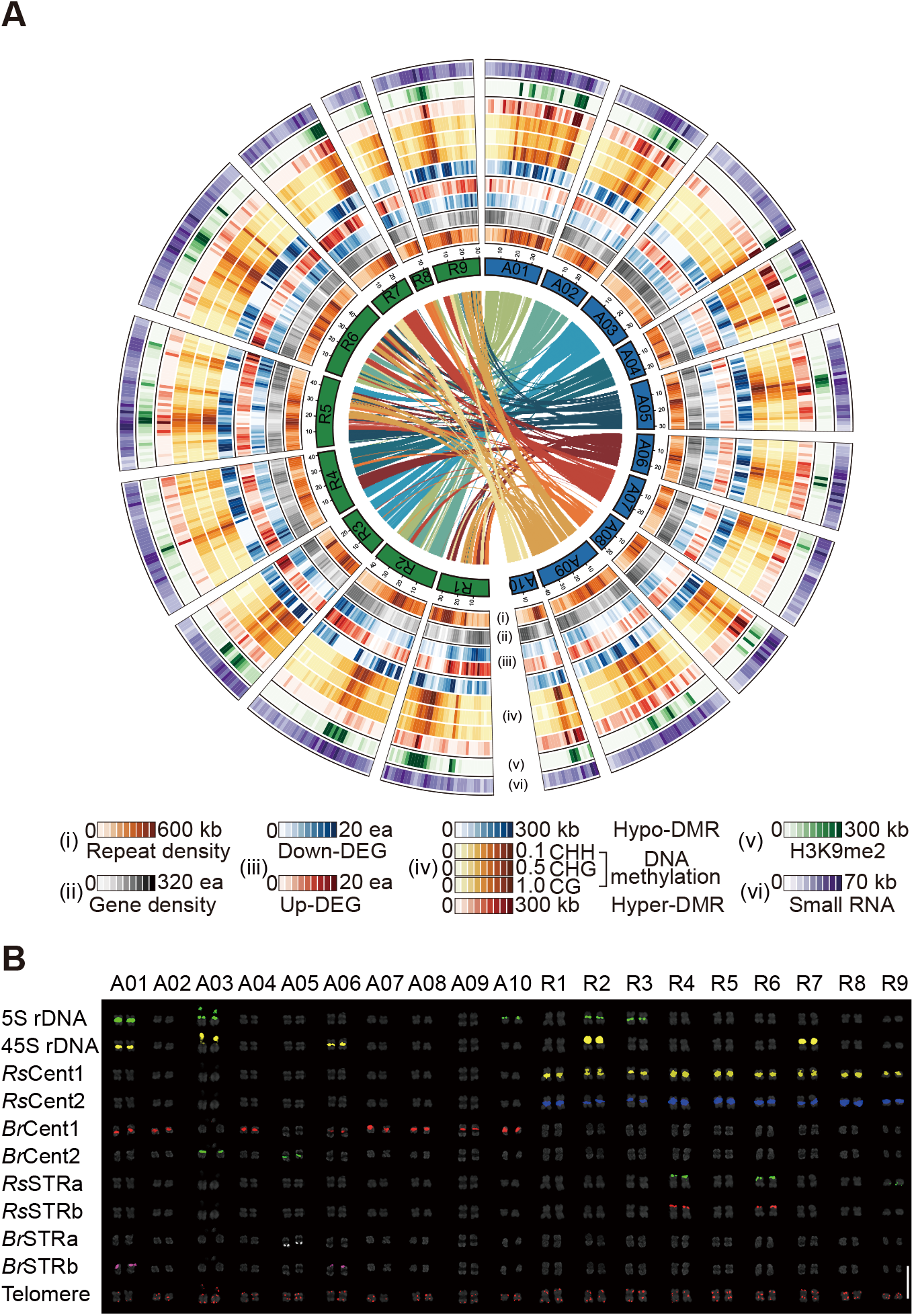
Genome structure of x*Brassicoraphanus*. A, The x*B* genome comprises 10 A_xB_ and 9 R_xB_ chromosomes. The data tracks represent (i) repeat density; (ii) gene density; (iii) DEGs between x*B* and its progenitor seedlings; (iv) CG, CHG, and CHH methylation levels and DMRs; (v) H3K9me2 repressive histone mark; and (vi) small RNAs. Lines in the inner circle represent syntenic relationships between A_xB_ and R_xB_. B, Multicolor Fluorescence *in situ* Hybridization (FISH) karyograms of x*B* with specific probes for 5S rDNA, 45S rDNA, centromeric tandem repeats (Cent), short tandem repeats (STR) and telomere repeats. Scale bars = 10 μm.

**Table 1.** Summary of the xBrassicoraphanus genome assembly.

| Assembly information |  | Contig | Scaffold |
| --- | --- | --- | --- |
| Total length / Number |  | 652.44 Mb / 68,454 ea | 692.83 Mb / 20,299 ea |
| Average / Median |  | 9.53 kb / 2.40 kb | 34.13 kb / 901 bp |
| Max / Min length |  | 190.62 kb / 200 bp | 16.46 Mb / 213 bp |
| N50 |  | 28,581 bp (6,854<sup>th</sup>) | 4,479,746 bp (49<sup>th</sup>) |
| N90 |  | 5,982 bp (24,969<sup>th</sup>) | 166,698 bp (284<sup>th</sup>) |
| GC contents |  | 35.75% | 33.68% |

| Scaffold assignment | Total number | Assigned to A<sub>xB</sub> genome | Assigned to R<sub>xB</sub> genome | Unassigned |
| --- | --- | --- | --- | --- |
| No. of scaffolds | 20,299 | 7,790 | 7,364 | 5,145 |
| Cumulative size (bp)<br>(% of total assembly) | 692,831,961<br>(100%) | 335,554,805<br>(48.43%) | 343,544,771<br>(49.59%) | 13,732,385<br>(1.98%) |
| No. of scaffolds assigned to reference chromosomes | 213 | 129 | 84 |  |
| Size of scaffolds assigned to reference chromosomes (bp)<br>(% of total assembly) | 581,691,615<br>(83.96%) | 279,795,674<br>(83.38%) | 301,895,941<br>(87.87%) |  |

| Species | Protein-coding loci | Total CDS length (bp) | Average CDS length (bp) | Average exon length (bp) | Average intron length (bp) |
| --- | --- | --- | --- | --- | --- |
| *xBrassicoraphanus* | 87,861 | 106,896,611 | 1,216 | 244 | 196 |
| *B. rapa* | 42,601 | 49,456,892 | 1,172 | 233 | 209 |
| *R. sativus* | 52,326 | 67,790,376 | 1,295 | 252 | 170 |

### Suppression of homoeologous interactions between A and R chromosomes

Interspecific hybridization often involves extensive homoeologous exchanges during meiosis eventually causing non-homologous recombination in immediate offspring (Szadkowski et al., 2010; Szadkowski et al., 2011; Xiong et al., 2011; Zhang et al., 2013; Grandont et al., 2014; Chen et al., 2018; Sosnowska et al., 2020). To investigate whether homoeologous interactions occur between A_xB_ and R_xB_ chromosomes, we examined the synapsis formation of meiotic chromosomes by immunolocalization of ASYNAPTIC1 (ASY1) and ZIPPER1 (ZYP1). ASY1 is the axial/lateral element of meiotic chromosomes loaded onto chromatids before synapsis (Armstrong et al., 2002), and ZYP1 is the central element of synaptonemal complex present in synapsed chromosomes (Higgins et al., 2005). We found that ASY1 was correctly loaded onto the entire axis of all euploid and allodiploid pachytene chromosomes at meiotic prophase I (Figure 2). ZYP1 also co-localized with ASY1 in all euploid pachytene chromosomes (Figure 2). Allodiploid *B. napus* (AC) produced discontinuous stretches of ZYP1 signals, indicating partial synapsis between A and C chromosomes (Supplemental Figure S7). Notably, however, ZYP1 was hardly associated with allodiploid x*B* (AR) pachytene chromosomes (Figure 2), suggesting that crossover between non-homologous chromosomes was strongly suppressed in x*B* (Park et al., 2020).

**Figure 2.**
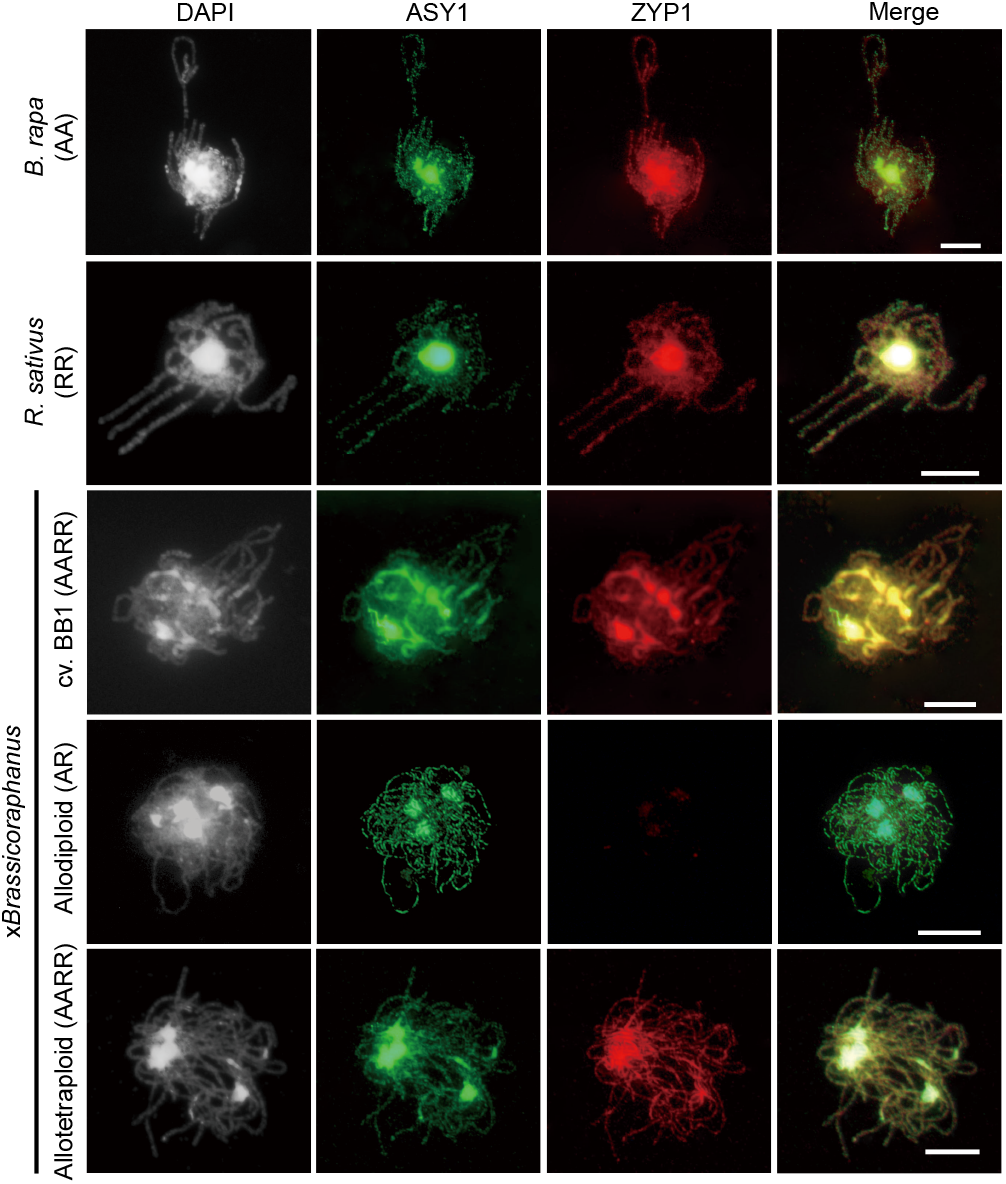
Chromosome behaviors of x*Brassicoraphanus*. Coimmunolocalization of ASY1 (green) and ZYP1 (red) at pachytene in *Br* (AA), *Rs* (RR), x*B* cv. BB1 (AARR), and resynthesized allodiploids (AR) and allotetraploid (AARR) x*B*. Chromosomes were stained with DAPI (white) and the overlay of three signals is shown (merge). Scale bars = 10 μm.

These findings demonstrate that *Br* and *Rs* chromosomes share little structrual similarity, and thus, orthology-dependent homoeologous interactions are prevented during meiosis while minimizing non-homologous exchanges, which would otherwise lead to aneuploidy and/or chromosome reshuffling. This also supports our observation that both *Br* and *Rs* genomes exist in entirety without losses in allotetraploid x*B* after hybridization (Figure 1B).

### Homoeologous expression adjustments in x*B*

It is assumed that speciation between *Br* and *Rs* has occurred earlier than between *Br* and *Bo*, although exact speciation timing is controversial (Mitsui et al., 2015; Jeong et al., 2016; Kim et al., 2018). Pairwise comparison of coding sequences (CDS) of all orthologs revealed 95.7% of sequence identity between *Br* and *Bo* within the same genus but 91.9% (*Br* vs. *Rs*) and 92.0% (*Bo* vs. *Rs*) across the genera (Figure 3, A and B). The same analysis in tribe Camelineae also showed similar sequence divergence for interspecific (93.5% for *A. thaliana* vs. *A. lyrata*) and intergeneric (89.7% for *A. lyrata* vs. *Capsella rubella*, and 90.3% for *A. thaliana* vs. *C. rubella*) relationships (Figure 3, A and B). Such divergence allowed us to clearly distinguish *Br*- and *Rs*-originated transcripts in x*B* (Figure 3, A and B). In x*B* seedling transcriptome, about half of the reads (51.4%) were assigned to A_xB_ and the other half to R_xB_ (48.6%), indicating that both subgenomes equally contribute to x*B* transcriptome (Supplemental Figure S8A). Similar portions of A_xB_ and R_xB_ transcripts were also present in four different tissues (leaf, hypocotyl, root and flower; Supplemental Figure S8A).

**Figure 3.**
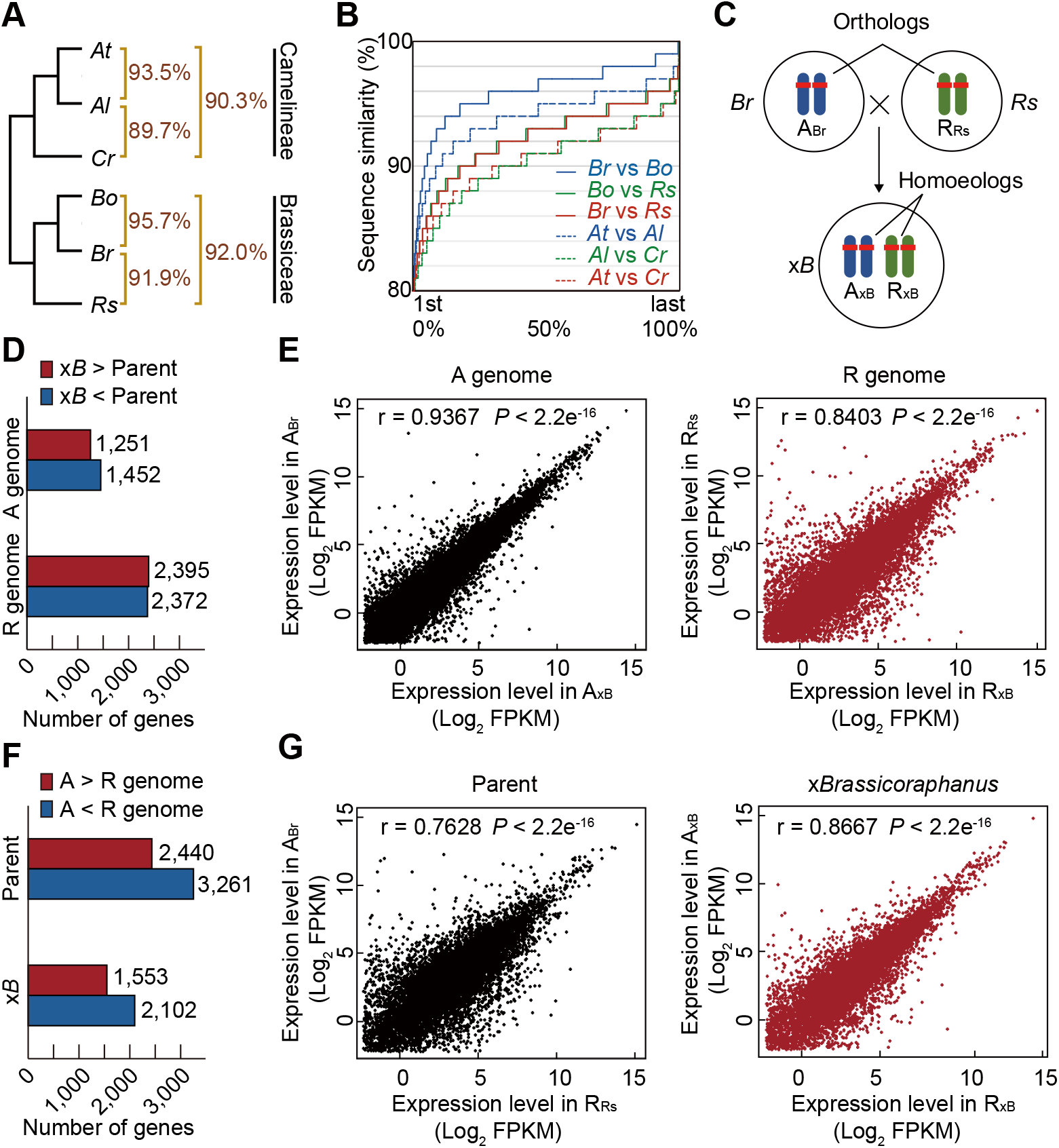
Transcriptome changes in x*B*. A, Phylogenetic relationship and sequence divergence in Camelineae and Brassiceae tribes. Percentages between species represent their CDS similarity of orthologous gene pairs. *At*, *Arabidopsis thaliana*; *Al*, *A. lyrata*; *Cr*, *Capsella rubella*. B, Distribution of sequence similarities of interspecific/intergeneric orthologs. Horizontal axis indicates orthologous gene pairs sorted in ascending order of sequence similarity. C, Relationship between orthologous and homoeologous genes in progenitors and x*B*. D, Number of DEGs in x*B* relative to the progenitors (A_Br_ vs. A_xB_ and R_Rs_ vs. R_xB_). E, Scatter plots comparing gene expression levels between A_Br_ and A_xB_ (black), and R_Rs_ and R_xB_ (red). F, Number of DEGs of orthologous pairs between A_Br_ and R_Rs_, and homoeologous pairs between A_xB_ and R_xB_. G, Scatter plots comparing gene expression levels between A_Br_ and R_Rs_ (black), and A_xB_ and R_xB_ (red).

Both *Br* and *Rs* genomes are retained, and thus, orthologous pairs become homoeologous each other in x*B* (Figure 3C). Among 28,751 genes commonly annotated in *Br* and x*B*, the majority were expressed at similar levels but 2,703 (9.40%) genes differentially expressed (>2 fold) between *Br* and x*B* seedlings (1,251 up-DEGs and 1,452 down-DEGs in x*B*; Figure 3, D and E). Differential expression between *Rs* and x*B* was more prominent, with 4,767 (20.96%) from 22,741 *Rs*-derived genes being dissimilarly expressed between *Rs* and x*B* seedlings (2,395 up-DEGs and 2,372 down-DEGs in x*B*; Figure 3, D and E). In addition, expression levels of *Br*-originated genes expressed in *Br* and x*B* seedlings were more positively correlated (r = 0.9367) than those of *Rs*-originated genes expressed in *Rs* and x*B* seedlings (r = 0.8403). These findings indicate that the majority of genes retain parental gene expression levels in x*B*, albeit *Br*-originated genes have a greater tendency to maintain their parental expression levels than *Rs*-originated genes. In other words, *Br* genome retains ‘maintenance expression’ over *Rs*, where *Br*-originated expression levels are preferentially inherited to the x*B* hybrid genome.

A total of 15,376 genes were identified as syntenic orthologs between *Br* and *Rs*, where 5,701 orthologous pairs (37.07%) were differentially expressed (>2 fold) between *Br* and *Rs* seedlings (2,440 up- and 3,261 down-DEGs in *Br* relative to *Rs*; Figure 3F). This indicates that *Br* and *Rs* have distinct expression profiles for phenotypic divergence. In x*B* seedlings, however, only 3,655 (23.77%) homoeologous pairs were differentially expressed (1,553 up- and 2,102 down-DEGs in A_xB_ relative to R_xB_; Figure 3F). Moreover, expression levels of A_xB_ and R_xB_ homoeologous pairs in x*B* seedlings were more highly correlated (r = 0.8667) than those of A_Br_ and R_Rs_ orthologous pairs between *Br* and *Rs* seedlings (r = 0.7628) (Figure 3G). This suggests that distinct expressions of many orthologous genes are adjusted to similar levels in the context of homoeologous relationship in x*B*. Such expression adjustment was also observed in tissue-specific expression profiles (Supplemental Figure S8B).

### Reconfiguration of transcription network

Previous studies analyzed the changes of expression levels with the sum of homoeologous pairs in allopolyploids relative to the parents, and determined additive or non-additive expressions of duplicated genes (Rapp et al., 2009; Grover et al., 2012; Yoo et al., 2014; Li et al., 2020; Shan et al., 2020; Wei et al., 2021). In this study, we further investigated how orthologous pairs were adapted to a new nuclear environment by monitoring changes of expression patterns of homoeologous genes in x*B* relative to the progenitors (Figure 4A). Out of 12,150 orthologous/homoeologous pairs commonly expressed in all *Br*, *Rs* and x*B* seedlings, 7,631 (62.80%) pairs were expressed at similar levels in every genome context, and their expressions are regarded to be ‘constant’ (gray in Figure 4A). By contrast, 1,435 (11.81%) pairs showed ‘biased’ expressions with significant differences between *Br* and *Rs*, while maintaining distinct progenitor expression levels in subgenomes A_xB_ and R_xB_ (blue in Figure 4A). Interestingly, expressions of 1,971 (16.22%) homoeologous pairs were adjusted to similar levels in x*B*, albeit their expressions were different between A_Br_ and R_Rs_ progenitors (red in Figure 4A). Such ‘convergent’ expressions were more prominent for R_Rs_-originated genes (1,483/1,971). We assumed that ‘convergent’ expressions might result from similar *cis*-regulatory sequences between homoeologous pairs under the same transcriptional control in x*B*. We analyzed the sequence similarities between homoeologous gene pairs of the categories of ‘convergent’ vs. ‘biased’ expressions (Figure 4B). Coding sequences of both ‘convergent’ and ‘biased’ homoeologous pairs have a high level of sequence identities (92.54% vs. 92.00%; Figure 4B). By contrast, the upstream *cis*- elements are noticeably divergent between homoeologous pairs. Interestingly, ‘convergent’ homoeologous pairs share less diverged *cis*-element sequences than ‘biased’ ones (68.42% vs. 63.29%; Figure 4B). These findings support our hypothesis that the upstream regulatory sequences of the orthologs have diverged after speciation but retain essential *cis*-elements that are likely under control of the same *trans*-acting regulators in x*B*. This also suggests that both A and R genomes still maintain the compatibility in transcription system to prevent a ‘transcriptome shock’ (Hegarty et al., 2006; Buggs et al., 2011), but divergence in regulatory elements should entail the reconfiguration of overall expression network in the hybrid genome of x*B*.

**Figure 4.**
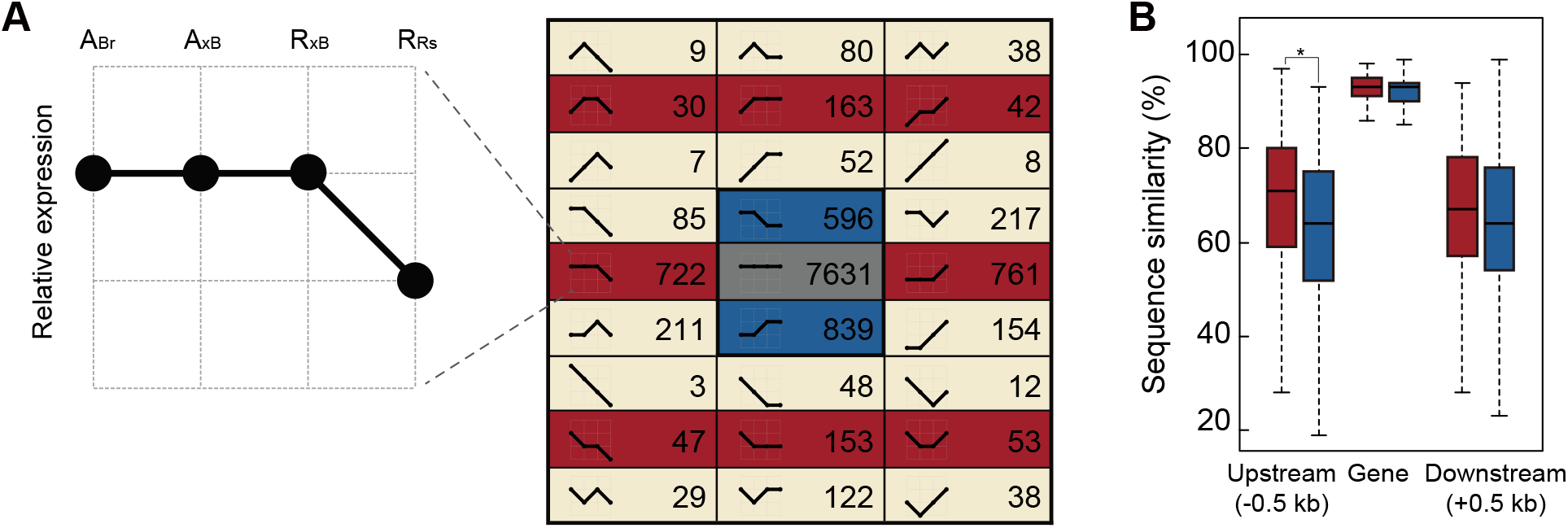
Expression patterns of homoeologous pairs in x*B*. A, Classifications of expression patterns of homoeologs in the x*B* relative to progenitor orthologs. The gray, blue and red blocks represent gene pairs showing ‘constant’, ‘biased’ and ‘convergent’ expressions, respectively. B, Sequence similarities of genic and adjacent upstream/downstream regions of orthologous genes showing convergent (red) and biased (blue) expressions in x*B* subgenomes (Wilcoxon’s rank-sum test, *P < 2.2e^-10^).

### Coordinated expression of homoeologous genes in response to external stimuli

Gene ontology (GO) enrichment analysis was performed for three categories of homoeologous expressions – ‘constant’, ‘biased’ and ‘convergent’. ‘Constant’ homoeologous pairs have enrichment for GO terms such as “cell differentiation”, “developmental cell growth” and “cell cycle” (Figure 5A and Supplemental Data Set S3), suggesting that cell function-related genes maintain consistent expression patterns after hybridization. However, the ‘biased’ homoeologous pairs did not display GO enrichment for specific functions (*P* > 0.001). Notably, the ‘convergent’ homoeologous pairs had GO enrichment for diverse responses such as “response to hormone”, “response to stress”, “response to biotic stimulus” and “response to abiotic stimulus” (Figure 5A and Supplemental Data Set 3). This suggests that the homoeologous pairs coordinately expressed in response to various stimuli tend to have similar *cis*-elements, although they are distinctly expressed in the progenitors.

**Figure 5.**
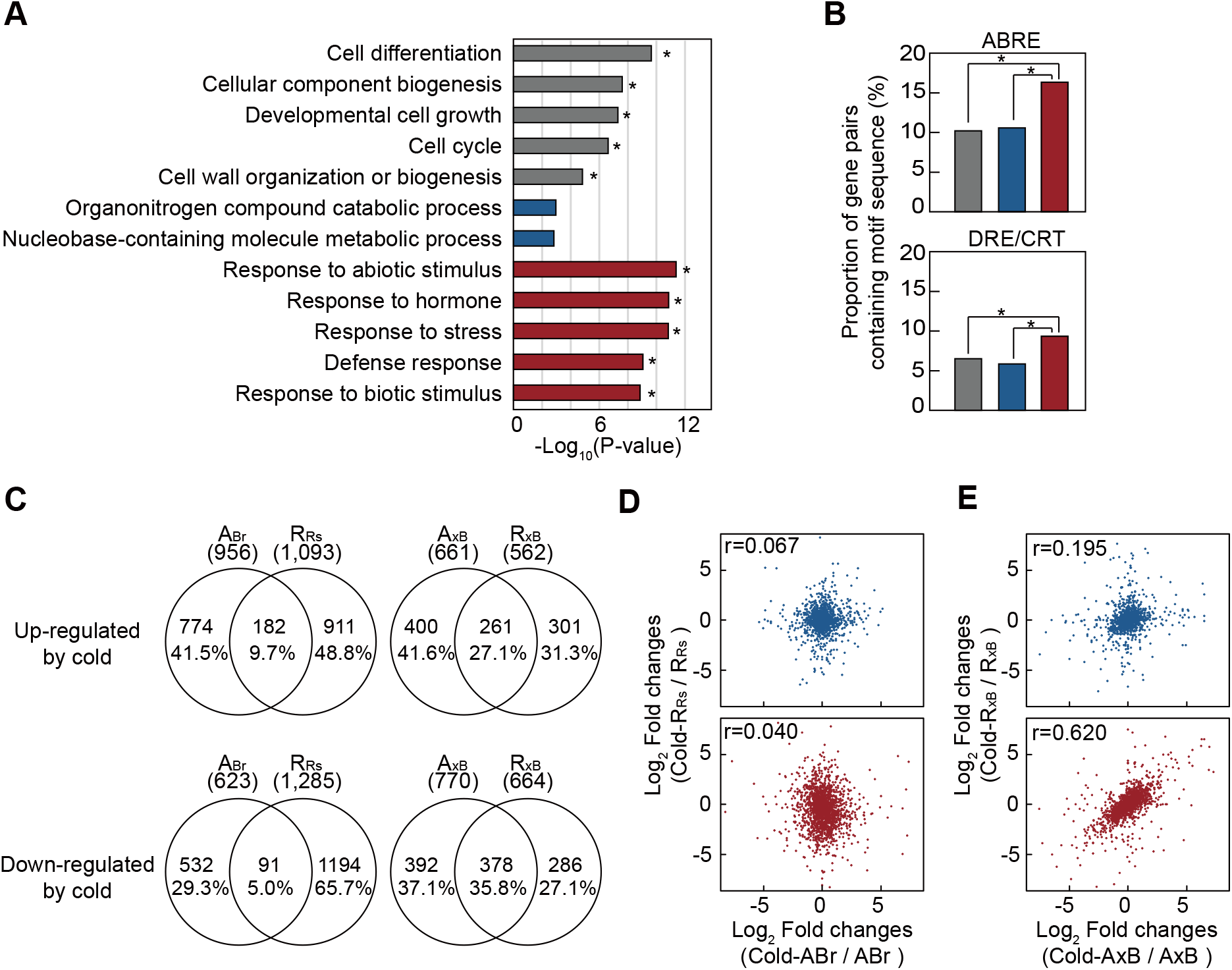
Expression of homoeologous genes in response to external stimuli. A, GO enrichments of ‘constant’ (gray), ‘biased’ (blue) and ‘convergent’ (red) homoeologous pairs (Fisher’s exact test, *P < 0.001). B, Proportion of ‘constant’ (gray), ‘biased’ (blue) and ‘convergent’ (red) homoeologous pairs containing conserved sequences of abscisic acid-responsive element (ABRE) and dehydration-responsive element/C-repeat element (DRE/CRT) (Fisher’s exact test, *P < 0.001). C, Venn diagram of cold-induced DEGs between A_Br_ and R_Rs_ orthologs (left) and between A_xB_ and R_xB_ homoeologs (right). D, Scatter plots of cold-induced expression changes of A_Br_ and R_Rs_ orthologous genes showing ‘biased’ (blue) and ‘convergent (red)’ expressions. E, Scatters plots of cold-induced expression changes of A_xB_ and R_xB_ homoeologous genes showing ‘biased’ (blue) and ‘convergent’ (red) expressions.

Moreover, the motifs of stress-responsive *cis*-elements such as abscisic acid-responsive element (ABRE; BACGTGK, B = C, G or T; K = G or T) (Lieberman-Lazarovich et al., 2019) and dehydration-responsive element/C-repeat element (DRE/CRT; RCCGAC, R = A or G) (Suzuki et al., 2005) were found abundantly in the upstream sequence of ‘convergent’ homoeologous pairs (Figure. 5B). This indicates that the genes involved in cellular signaling may require essential *cis*-elements to properly respond to external stimuli.

We treated cold to *Br*, *Rs* and x*B* seedlings and monitored expression changes of orthologous/homoeologous genes. Out of 15,376 orthologs, 1,579 genes were differentially regulated by cold in *Br* seedlings, with 956 up-DEGs and 623 down-DEGs (Figure 5C). In cold-treated *Rs* seedlings, 2,378 genes were differentially expressed, with 1,093 up-DEGs and 1,285 down-DEGs (Figure 5C). Among them, only small fractions of orthologous genes (182 up- and 91 down-DEGs; 9.75% and 5.01%) were similarly regulated in both *Br* and *Rs*(Figure 5C). In x*B* seedlings, a total of 2,657 genes were differentially regulated by cold treatment. Specifically, 1,431 *Br*-derived orthologs were differentially expressed (661 up-DEGs and 770 down-DEGs in A_xB_) and 1,226 *Rs*-derived orthologs differently regulated (562 up-DEGs and 664 down-DEGs in R_xB_) after cold treatment (Figure 5C). Notably, a larger fraction (261 up- and 378 down-DEGs; 27.13% and 35.80%) of A_xB_ and R_xB_ homoeologous pairs were identified as common DEGs in x*B* (Figure 5C). These observations indicate that many of orthologous/homoeologous pairs are distinctly regulated in *Br* and *Rs* progenitors but their expressions are systematically coordinated in x*B* hybrid genome in response to cold exposure. We also found that expressions of A_Br_ and R_Rs_ orthologous genes had a weak correlation regardless of expression categories (Figure 5D). Interestingly, A_xB_ and R_xB_ ‘convergent’ homoeologous pairs had a strong correlation (r = 0.620), whereas ‘biased’ ones did not (r = 0.195) (Figure 5E). These data suggest that evolutionarily divergent homoeologous pairs still share essential motifs in *cis*-elements that can be subjected to the same *trans*-acting regulation, conceivably responsible for coordinated expressions in response to environmental cues in hybrids.

### Silencing of transposable element stabilizes the x*B* hybrid genome

Resynthesized hybrids often experience epigenetic alterations (Greaves et al., 2015). We investigated methylation profiles in coding genes and repeat regions. In coding regions, DNA methylation levels are high in gene body, decrease towards 5’ and 3’ shores, and increase again beyond translation start and termination sites in all *Br*, *Rs* and x*B* seedlings (Figure 6A). Notably, A_Br_ and R_Rs_ progenitor genomes have distinct CG methylation patterns in coding genes, with A_Br_ being more densely methylated than R_Rs_. This methylation asymmetry is inherited to A_xB_ and R_xB_ subgenomes (Figure 6A). TEs are heavily methylated in general, especially near-complete CG methylation in all species (Figure 6B). TEs also have higher CHG and CHH methylation levels than coding genes. Interestingly, *Br* and *Rs* TEs have distinct CHG methylation profiles, with more CHG methylation at *Rs* TEs (Figure 6B). However, such asymmetry is abolished in x*B*, where *Br*-derived TEs have an increased CHG methylation level comparable to *Rs*-derived TEs (Figure 6B). This suggests that TEs from *Br* acquired more CHG methylation after hybridization possibly via *trans*-acting mechanisms.

**Figure 6.**
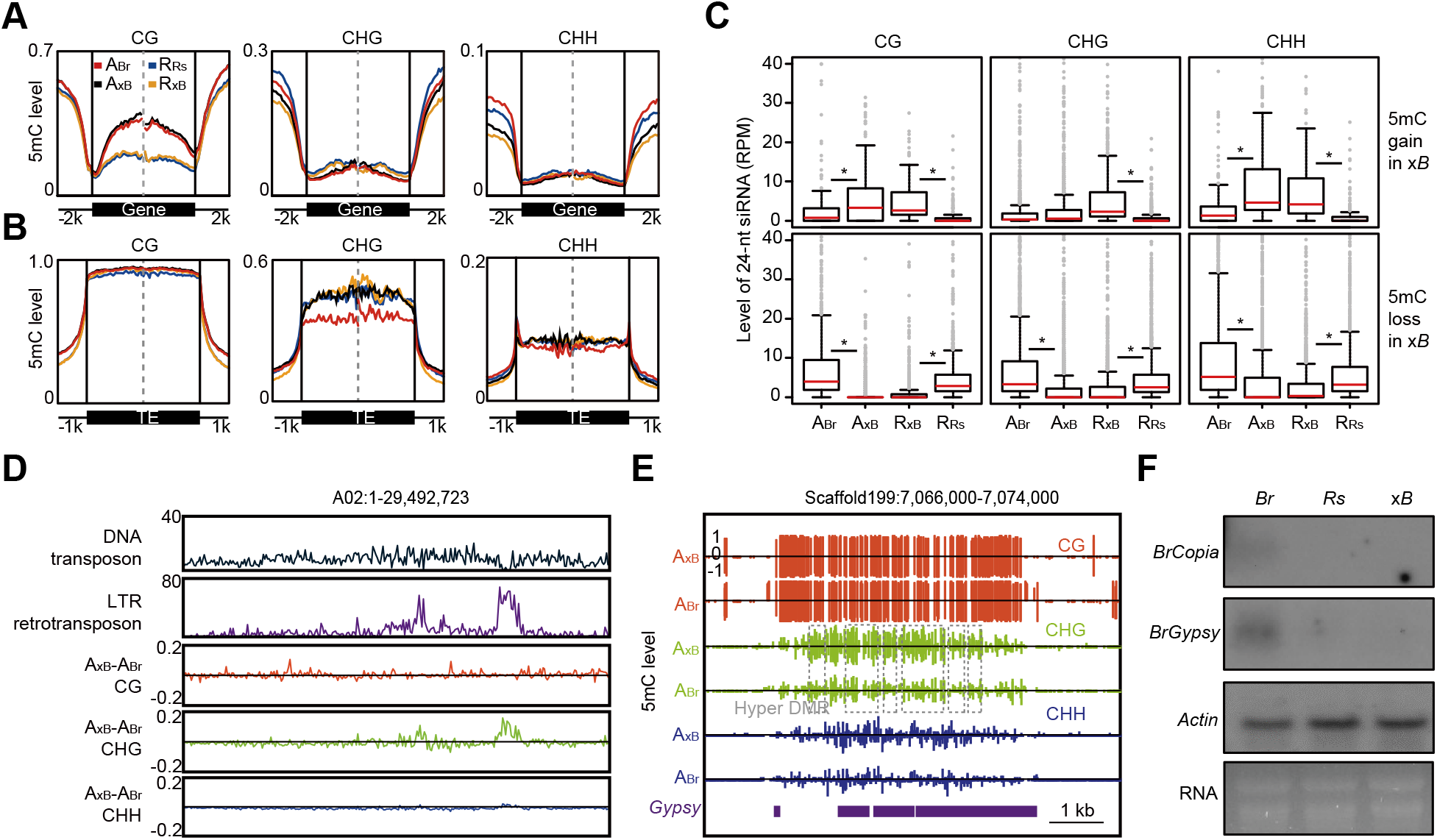
Relationships of DNA methylation, small RNA and TE expression in x*Brassicoraphanus*. A and B, Distribution of DNA methylation at gene body (A) and TE regions (B) in x*B* subgenomes (A_xB_ and R_xB_) and its progenitor genomes (A_Br_ and R_Rs_). C, Expression levels of 24-nt RNAs at CG, CHG and CHH DMRs in x*B* subgenomes (A_xB_ and R_xB_) and the progenitor genomes (A_Br_ and R_Rs_). The expression level of 24-nt RNAs was calculated as reads per million (RPM) (two-tailed Student’s t-test, *P < 5.0e^-5^). D, Distributions of DNA transposons, LTRs and DNA methylation difference between A_Br_ and A_xB_ across chromosome A02 in 100 kb bins. E, An example of methylation distributions at hypermethylated *Gypsy* class LTR in A_xB_ and A_Br_. F, Northern blot for *BrCopia* and *BrGypsy*. Actin was used as a loading control.

We analyzed small RNAs in *Br*, *Rs* and x*B* seedlings, and found that approximately 30∼50% of small RNAs were 24-nt RNAs as potential short-interfering RNAs (siRNAs) (Supplemental Figure S9A). siRNAs were highly associated with hyper-DMRs in x*B* but loosely with hypo-DMRs, indicating a strong correlation between 24-nt RNA and DNA methylation (Figure 6C). About 12% of 24-nt RNAs from *Br* and *Rs* have a pairwise sequence identity and may share the same targets across the genomes (Supplemental Figure S9B). Indeed, 10.4% of 24-nt RNAs from x*B* also have indistinguishable origins (Supplemental Figure S9C). This suggests that, in x*B* hybrid genome, R_xB_-originated siRNAs induce gain of CHG methylation at TEs on A_xB_ possibly via RNA-directed DNA methylation (RdDM) (Law and Jacobsen, 2010). DNA transposons are widespread throughout the x*B* genome with little association with DMRs (Figure 6D). LTRs that account for approximately 30% of repeats (Supplemental Table S3) were also heavily methylated. Notably, it was clear that LTRs on A_xB_ had higher methylation levels at the CHG context than A_Br_ (Figure 6D and, Supplemental Figures S10 and S11). This suggests that DNA methylation profiles have changed in a subgenome-specific manner, for which R_xB_-originated siRNAs might induce gain of CHG methylation in *trans* at LTRs of the same kind on A_xB_. As exemplified in Figure 6E, the *Gypsy* element on A_xB_ was found to have higher CHG methylation levels than A_Br_ at the scaffold level, albeit CG and CHH methylation levels are nearly identical. Northern blot analysis verified that *Copia* and *Gypsy* elements were moderately expressed in *Br* but silenced in x*B* seedlings (Figure 6F). These findings suggest that RdDM-mediated DNA methylation induces TE silencing across subgenomes, which in turn stabilizes the x*B* hybrid genome.

## Discussion

Hybridization barriers serve as a mechanism to prevent a gene flow between species (Abbott et al., 2013). In particular, the post-zygotic hybridization barrier after fertilization is often manifested as hybrid inviability or sterility (Dion-Cote and Barbash, 2017). Hybrid sterility is generally associated with a failure in meiosis. Normal meiosis requires the formation of synapsis between homologous chromosome pairs, but when they are abolished or formed between multiple and/or non-homologous chromosomes, the chromosomes segregate abnormally, resulting in sterile gametes and aneuploidy (Martinez-Perez and Colaiacovo, 2009). Aneuploidy and/or chromosome rearrangements are frequently observed in resynthesized allopolyploids between close species (Xiong et al., 2011; Zhang et al., 2013; Chen et al., 2018). This is mainly caused by the collinearity/homology between less divergent parental chromosomes. For instance, A1/C1, A2/C2 and parts of A5/C4 (A from *Br* and C from *Bo*) chromosomes are homologous to each other (Parkin et al., 2005), and most phenotypic variations and aneuploidy in resynthesized *B. napus* lines are caused by homoeologous interactions, mostly between non-homologous chromosomes (Gaeta et al., 2007; Xiong et al., 2011; Grandont et al., 2014). However, the presence of full compliments of both *Br* and *Rs* chromosomes in x*B* demonstrates that a merger of divergent genomes may avoid such harmful interactions, while producing fertile gametes after polyploidization.

Hybridization between species inevitably entails changes in *cis*-*trans* interactions bringing about alterations in the transcription network (Hu and Wendel, 2019). Therefore, extensive changes in parental expression profiles are expected, and when such changes are intolerable, the hybrid will undergo a ‘transcriptome shock’, manifested as hybrid dysgenesis (Martienssen, 2010) or outcrossing depression (Frankham et al., 2011). x*B* experienced moderate expression changes of progenitor genes after hybridization but still maintains a transcription network between subgenomes compatible enough to generate novel or intermediate phenotypes. Our four-point expression analysis revealed that ‘convergent’ homoeologs share similar *cis*-elements, and expression levels of a larger fraction of *Rs*- derived homoeologs were adjusted to *Br*-derived ones. This suggests that *Br*-originated *trans*- acting factors probably play dominant roles for co-regulation of homoeologous pairs in x*B* (Hu and Wendel, 2019). Notably, stress response-related motifs are enriched in the *cis*- elements of ‘convergent’ homoeologs, suggesting that transcriptional regulation is primarily mediated by *trans*-acting factors sharing common homoeologous targets that are involved in diverse responses. Such reconfiguration of transcription network is conceivably crucial to the adaptation of newly synthesized hybrids.

*Br* has higher gene-body CG methylation levels than *Rs*, which is inherited to each subgenome in x*B*. This indicates that differential gene-body methylation is maintained after hybridization and this methylation asymmetry may contribute to ‘maintenance expression’ of A_xB_ through unknown mechanisms. TEs are heavily methylated in general, but also showed asymmetric CHG methylation between *Br* and *Rs*. Intriguingly, *Br*-originated LTRs gained CHG methylation comparable to *Rs* ones in x*B*, suggesting that repeat-originated siRNAs trigger hypermethylation via RdDM in *trans* and TE silencing (Wendel et al., 2016). This may prevent hyper-activation of TEs and subsequent genome destabilization, which would otherwise culminate to a ‘genomic shock’ as initially proposed by McClintock (McClintock, 1984).

It is believed that the more distantly related the species, the stronger the hybridization barrier. On the contrary to this assumption, our findings strongly suggest that, as long as the physiology and transcriptional regulatory networks are compatible, a certain extent of genome divergence promotes hybridization between distant species. Therefore, a trade-off between genome divergence and transcriptome compatibility is meaningful to facilitate hybridization between species without causing genome destabilization and/or a conflict in transcription network. This concept also proposes that interspecific/intergeneric hybridization may occur more frequently in nature than we have thought, and the model of ‘triangle of U’ (U, 1935) can be further expanded to the intergeneric level.

After whole genome duplication or hybridization between the different species followed by chromosome doubling (allopolyploidization), polyploid plants generally undergo gradual but substantial genome reconstruction including massive chromosome rearrangement, differential deletion or retention of duplicated genes and biased genome fragmentation (Cheng et al., 2018). This eventually leads to a decrease in both chromosome number and genome size, with most of polyploid properties being lost. Extensive changes in genome structure and gene repertoire accompanied with sub-functionalization/neo- functionalization of duplicated genes also contribute to the formation of new species with novel phenotype and function, which sometimes outperform the diploid progenitors with the greater ecological fitness. Thus, evolution of land plants, especially the angiosperms, is not a one-way process. Rather, it is likely to comprise the recurrent cycles of hybridization, diversification, diploidization and reunification among the species in the same lineage (Wendel, 2015). Furthermore, understanding the highly dynamic and flexible process of hybridization and polyploidization should provide a clue to Charles Darwin’s ‘abominable mystery’ (Darwin, 1903) questioning the great diversification and expansion of angiosperms within a short geological time.

## Materials and methods

### Plant materials

x*Brassicoraphanus* cv. BB1 (x*B*), *Brassica rapa* L. cv. Chiifu-401-42 (*Br*), and *Raphanus sativus* L. cv. WK10039 (*Rs*) were grown on 1x Murashige and Skoog (MS) medium (Duchefa) in a growth chamber under 16 hr of fluorescent light at 20 ± 10 μmol m^-2^ s^-1^, 22°C for 14 days. The seedlings including shoots and roots were harvested together for whole genome-seq, RNA-seq, bisulfite (BS)-seq, chromatin immunoprecipitation (ChIP)-seq and small RNA-seq. For tissue-specific transcriptome analysis, RNA was extracted from leaf, hypocotyl, and root of the seedling and from the opened flower of *Br*, *Rs* and x*B*. For cold treatment, 14-day-after-sowing seedlings of *Br*, *Rs* and x*B* were grown at 4°C for five weeks. RNA was extracted and stored at -20°C until use.

### Genome sequencing, assembly and genome size estimation

Paired-end and mate-pair sequencing libraries with insert sizes of 200 bp, 400 bp, 3 kb, 8 kb 5 kb, 10 kb and 15 kb were constructed using KAPA library prep kit (Roche) and Illumina Mate Pair Library kit (Illumina) following the manufacturer’s instructions (Supplemental Table S1). The libraries were sequenced on an Illumina HiSeq 2000 platform. Prokaryotic sequences, duplicated reads, low quality reads and low frequency reads were filtered out (Supplemental Table S1). The preprocessed sequences were assembled using SOAPdenovo2 (Luo et al., 2015) with the best *k*-mer values for each library. To increase the length of scaffolds, serial scaffolding processes were carried out using SOAPdenovo2 (Luo et al., 2015) and SSPACE (Boetzer et al., 2011). Gaps in the scaffolds were reduced further using SOAPdenovo Gapcloser (Luo et al., 2015) and Platanus (Kajitani et al., 2014) (Supplemental Table S2). In the k-mer analysis, counting k-mer occurrence of 19-mer were performed using Jellyfish (Marcais and Kingsford, 2011). The genome size of x*B* was estimated by flow cytometry analysis (FACSCalibur, BD Biosciences) as previously described (Huang et al., 2013). Genome data were visualized with Circos (Krzywinski et al., 2009).

### Chloroplast genome assembly

The chloroplast genome was *de novo* assembled from the 1x coverage of whole-genome sequencing reads. The chloroplast genome was annotated with GeSeq (Tillich et al., 2017) and manually curated. The chloroplast genome was visualized using OrganellarGenomeDRAW (Lohse et al., 2013).

### Assignment of scaffolds to A_xB_ and R_xB_ subgenomes

Whole-genome sequencing reads of *Rs* and *Br* from Brassica Database (BRAD) were mapped to the x*B* scaffolds using Bowtie (Langmead et al., 2009). The number of mapped reads was counted and the scaffolds were assigned to A_xB_ and R_xB_ subgenomes, based on a comparison of the number of parental reads (A_xB_ subgenome: >99% ratio of mapped reads from *Br*; R_xB_ subgenome: >99% ratio of mapped reads from *Rs*). Next, assigned x*B* scaffolds were anchored to the reference chromosomes of *Br* and *Rs* to build x*B* pseudo-chromosomes.

### Gene and TE annotation

Gene annotations of x*B* and *Rs* were performed following the previous annotation pipeline with minor modifications (Kim et al., 2014). Briefly, the annotation pipeline consisted of repeat masking, mapping of different protein sequence sets and mapping of RNA-seq reads. Independent *ab initio* predictions were performed with AUGUSTUS (Stanke et al., 2008). The EVidenceModeler (Haas et al., 2008) software combines *ab initio* gene predictions with protein and transcript alignments into weighted consensus gene structures. Gene annotation of *Br* was downloaded from Ensembl plant (ftp://ftp.ensemblgenomes.org/pub/plants/release-31/gff3/brassica_rapa/) and additional 1,700 genes were annotated using Exonerate (Slater and Birney, 2005). Functional annotation was performed through BLASTP against SwissProt and Plant RefSeq database. TE-related repeat sequences were predicted by RepeatModeler (Smit and Hubley, 2008) and Repeatmasker (Smit et al., 2015).

### Fluorescence *in situ* hybridization (FISH) analysis

The sequences of 5S rDNA, 45S rDNA, *Rs*Cent1, *Rs*Cent2, *Br*Cent1, *Br*Cent2, *Rs*STRa, *Rs*STRb, *Br*STRa, *Br*STRb and telomere were used as probes (Supplemental Table S5). The probes were labelled by nick translation with different fluorochromes. Root mitotic chromosome spreads and FISH procedures were performed according to the previous method (Waminal and Kim, 2012). For directly labelled probes, slides were immediately used for FISH after fixation with 4% paraformaldehyde, without subsequent pepsin and RNase pretreatment. Images were captured with an Olympus BX53 fluorescence microscope equipped with a Leica DFC365 FS CCD camera and processed using Cytovision ver. 7.2 (Leica Microsystems).

### Resynthesized allodiploid and allotetraploid x*Brassicoraphanus* plants

Resynthesized allodiploid x*Brassicoraphanus* plants were produced from a cross between *B. rapa* cv. Chiifu-401-42 as the seed parent and *R. sativus* cv. WK10039 as the pollen donor. Thirty-day-old immature hybrid ovules were cultured on 1 × MS medium supplemented with 2% sucrose (w/v) and 0.8% plant agar (w/v). The plates were placed at 24°C growth chamber for two weeks and then seedlings were vernalized at 4°C cold chamber for 4 weeks with 16 hr of light and 8 hr of dark.

### Resynthesized allodiploid and allotetraploid *B. napus* plants

Resynthesized allodiploid *B. napus* plants were produced from a cross between *B. rapa* cv. Chiifu-401-42 as the seed parent and *B. oleracea* var. Capitata as the pollen donor. Ovary culture was performed as described in the published protocol with modifications (Inomata, 1977). Ovaries at 4 day after pollination were explanted on 1 × MS medium supplemented with 5% sucrose (w/v), 300 mg·L^-1^ casein hydrolysate and 0.8% plant agar (w/v) at 24°C growth chamber. Four weeks after explantation, hybrid ovules were transferred on B5 medium with vitamin supplemented with 2% sucrose (w/v), 300 mg·L^-1^ casein hydrolysate and 0.8% plant agar (w/v). Ovules were incubated in the dark at 24°C in three days and placed at 16 hr of light and 8 hr of dark condition. Seedlings were vernalized at 4°C cold chamber for 4 weeks with 16 hr of light and 8 hr of dark. The plants were transferred to pots in the greenhouse with the same light condition. A 0.3% colchicine solution was applied to the shoot apical meristems for two days to obtain allotetraploids.

### Production of antibody and immunolocalization of meiotic proteins

The coding regions of *BrASY1* and *BrZYP1* genes were PCR-amplified from cDNA of young flowering buds from *Br* (Supplemental Table S6). The fragments of *BrASY1* (708 bp) and *BrZYP1* (1,332 bp) were inserted into the pET-28a expression vector (Novagen) and transformed into *Escherichia coli* Rosetta2 (DE3) strains (Novagen). The transformed *E. coli* cells were grown at 30°C in 1 L of Luria-Bertani broth (LB) medium in the presence of 50 µg·mL^-1^ of kanamycin and 50 µg·mL^-1^ of chloramphenicol until the OD600 reached to 0.4.

Recombinant protein expression was induced with 1 mM of isopropyl b-D-thiogalactopyranoside (IPTG) at 16°C for 16 hr. Cells were centrifuged (4°C) at 6,500 rpm for 15 min and the pellet was resuspended in 100 mL of ice-cold column buffer (50 mM Tris-HCl, pH 7.4, 100 mM NaCl, 10% glycerol, 0.1 mM dithiothreitol, 0.1 mM PMSF). Cells were lysed by sonication for 5 min on ice (output power, 4; duty cycle, 50%; Branson Sonifer 250, Branson). Inclusion bodies were collected by centrifugation (4°C) at 9,000 rpm for 25 min and dissolved in 4 M urea. The soluble lysate was purified with a 5-mL HisTrap FF column (GE Health care, USA) with a linear gradient of ice-cold column buffer (50 mM Tris-HCl, pH 7.4, 100 mM NaCl, 10% glycerol, 0.1 mM dithiothreitol, 250 mM imidazole). The purified BrASY1 and BrZYP1 proteins were used to produce polyclonal antibodies from rabbit and rat, respectively, by Youngin Frontier (Korea), and the quality of antibody was validated by western blot. Immunolocalization was performed as described in the published protocol (Chelysheva et al., 2013). In brief, primary antibodies anti-BrASY1 and anti- BrZYP1 were used at dilution 1:250 in PBST (0.1% Triton-X 100 in 1× PBS) containing 1% BSA and the secondary antibodies (Goat anti-rabbit IgG H&L, Alexa Fluor 488 and Donkey anti-rat IgG H&L, Alexa Fluor 594) were used at dilution 1:500. Images were captured with an Axioskop2 microscope equipped with an Axiocam 506 color CCD camera (Zeiss) and processed using Adobe Photoshop CS6 (Adobe Systems Incorporated).

### Identification of orthologous and homoeologous gene pairs

To identify orthologous gene pairs between parental genomes (A_Br_ vs. R_Rs_), the reciprocal best BLAST hit was performed with >80% of identity and >80% of coverage. Syntenic regions were defined as contiguous regions containing at least five homologous gene pairs in *Br* and *Rs* genomes, and the pairs in the syntenic regions were determined as orthologous gene pairs. Homoeologous gene pairs between the progenitor genomes (A_xB_ vs. R_xB_) were determined following the same standard.

### RNA-seq analysis

Total RNA was extracted with RNeasy plant kit (Qiagen) following the manufacture’s protocol. The DNase-treated RNA samples, including two replicates for each of seedling, leaf, hypocotyl and flower, and one replicate for root of x*B* and its progenitors, were used for constructing RNA-seq libraries (Zhong et al., 2011). RNA sequencing was performed on an Illumina HiSeq 2000 platform. The obtained raw reads were filtered using FASTX-Toolkit and low quality reads (Q < 20) were removed. The filtered reads were mapped on *Br*, *Rs* and x*B* genomes using Tophat (Trapnell et al., 2009) with default parameters (Supplemental Data Set S4). The mapped read counts were calculated using HTSeq (Anders et al., 2015).

Statistical tests of DEGs were performed using EdgeR (Robinson et al., 2010) with the false discovery rate (FDR) < 0.05 and fragments per kilobase of transcript per million mapped reads (FPKM) log_2_ fold change > 1. The Gene ontology (GO) terms of x*B* genome were annotated by Blast2Go using the non-redundant sequence database from NCBI with < 1e^-15^ of e-value parameter. The statistical comparison of GO term accumulation was conducted using TopGo in R package (Alexa and Rahnenfuhrer, 2010) with p-values of fisher’s exact test (*P* < 0.001). Motifs of ABRE (BACGTGK, B = C, G or T; K = G or T) and DRE/CRT(RCCGAC, R = A or G) were searched in 500 bp upstream regions of genes using FIMO (Grant et al., 2011) with parameters “--verbosity 1 --thresh 0.01”.

### BS-seq analysis

Genomic DNA (5 μg) was used to construct the BS-seq library with the KAPA Library kit (Roche) and EpiTect Bisulfite Kit (Qiagen) according to manufacturer’s instructions. The libraries were sequenced using the Illumina HiSeq 2000. Raw reads were filtered using FASTX-Toolkit and low quality reads (Q < 20) were removed. Reads were mapped onto the x*B* genome using BISON (Ryan and Ehninger, 2014), with the parameters “--very-sensitive -- score-min ‘L,-0.6,-0.6’”. Only cytosine sites with 4x coverage read depths were accepted for the subsequent analysis. Differentially methylated cytosines (DMCs) and regions (DMRs) were identified as described previously (Kim et al., 2019). In brief, DMCs were identified using Fisher’s exact test (*P* < 0.05) between the levels of methylation in x*B* and the progenitors *Br* and *Rs*. DMRs were identified based on the regions with a length ≥ 200 bp, ≥ 5 DMCs, and the mean methylation difference ≥ 0.3 for CG, ≥ 0.15 for CHG, and ≥ 0.1 for CHH (Supplemental Data Set S1). For metagene plot of DNA methylation in gene bodies and repeat, regions of 2 kb upstream, downstream and gene body were divided into 50 bp windows and methylation levels were calculated each. Methylation data were visualized with the Integrated Genome Browser (Freese et al., 2016).

### ChIP-seq analysis

ChIP was performed following the published protocol (Lee et al., 2007). Chromatin was immunoprecipitated with antibody against histone H3K9me2 (Abcam). ChIP-seq libraries were constructed as described in the Illumina ChIP sequencing kit (Illumina). DNA fragments with about 600 bp were excised from an agarose gel and amplified for cluster generation and sequencing. All DNA libraries were sequenced on a HiSeq2500 platform (Illumina) with single-end reads. The sequencing reads were quality-controlled with FASTX-Toolkit and aligned to x*B* genome using Bowtie (Langmead et al., 2009) with parameters “- best -m1”. H3K9me2-enriched regions were defined using SICER (Zang et al., 2009) (window size = 500, gap size = 600, FDR = 0.01) and overlapping regions between two biological replicates were identified using the MergePeaks module of the Homer software (Heinz et al., 2010) (Supplemental Data Set S2).

### Small RNA-seq analysis

The small RNA libraries were constructed using the Illumina TruSeq Small RNA sample Prep kit (Illumina). The libraries were sequenced on the HiSeq 2000 platform (Illumina). The adaptor sequences were trimmed using cutadapt (Martin, 2011) with parameters “-g TACAGTCCGACGATC -a TGGAATTCTCGGGTGCCAAGG -m 18 -M 30”. Low quality sequences were removed using FASTX-Toolkit with parameters “-q 20 -p 100”. The quality-trimmed read sequences ranged from 18 to 30-nt were mapped to the x*B* genome using Bowtie (Langmead et al., 2009) with parameters “-best -v 0”. Mapped reads were classified into ribosomal RNA, small nucleolar RNA, small nuclear RNA, signal recognition particle RNA, and transfer RNA using Rfam database version 12.1 (Nawrocki et al., 2014).

Prediction of microRNA (miRNA) was performed with the miRDeep-P (Yang and Li, 2011) and ShortStack (Axtell, 2013), and the secondary structure was predicted using RNAfold.

Candidate miRNAs were annotated by alignment to miRBase database version 21 (Kozomara and Griffiths-Jones, 2013).

### Northern blot analysis

Total RNA (10 µg) was electrophoresed on a 1% formaldehyde denaturing gel and transferred onto the Hybond N+ membrane (GE Healthcare). The *BrGypsy*, *BrCopia* and *Actin* probes were amplified by PCR, and randomly labelled with [α-32P]dCTP (Perkin Elmer) using a Klenow fragment (3′ → 5′ exo−) (New England Biolabs). Hybridization was performed at 65°C overnight in the pre-hybridization solution containing 6x saline-sodium citrate buffer, 5x Denhardt’s reagent, and 1% sodium dodecyl sulphate. After hybridization, the membrane was washed and exposed to an X-ray film (Fujifilm). Primer sequences are provided in Supplemental Table S7.

### Accession numbers

The sequencing data for genomic, transcriptomic and epigenomic analyses are available from Bioproject ID PRJNA353741, PRJNA353738, PRJNA394950 and PRJNA353316. The assembled x*Brassicoraphanus* genome is available from Bioproject ID PRJNA353741. The chloroplast genome of x*B* is deposited to GenBank under accession number MN928713.

## Supplemental data

Supplemental Figure S1. Phenotypes of x*Brassicoraphanus* intermediate between *B. rapa* and *R. sativus*.

Supplemental Figure S2. Flow cytometry analysis and genome size estimation of x*Brassicoraphanus*.

Supplemental Figure S3. Chloroplast genome of x*Brassicoraphanus*.

Supplemental Figure S4. Comparison of x*Brassicoraphanus* genome with its parental genomes.

Supplemental Figure S5. Genome-wide DNA methylation in x*Brassicoraphanus*.

Supplemental Figure S6. H3K9me2 modification of x*Brassicoraphanus*.

Supplemental Figure S7. Chromosome interactions in resynthesized x*Brassicoraphanus* and *Brassica napus*.

Supplemental Figure S8. Transcriptome analysis of x*Brassicoraphanus*.

Supplemental Figure S9. Small RNA analysis of x*Brassicoraphanus*.

Supplemental Figure S10. DNA methylation metaplots in transposable elements.

Supplemental Figure S11. Distribution of DNA methylation changes in A and R subgenomes of x*Brassicoraphanus*.

Supplemental Table S1. Summary of genomic reads from x*Brassicoraphanus*.

Supplemental Table S2. Statistics of x*Brassicoraphanus* genome assembly.

Supplemental Table S3. Annotation of repeat sequences in x*Brassicoraphanus* genome.

Supplemental Table S4. Chloroplast genome annotations of x*Brassicoraphanus* and progenitors.

Supplemental Table S5. Primers and oligo for FISH probes.

Supplemental Table S6. Primers for production of antibody.

Supplemental Table S7. Primers for northern blot probes.

Supplemental Data Set S1. Detected differentially methylated regions in x*Brassicoraphanus*.

Supplemental Data Set S2. Histone H3K9me2 peak regions in *B. rapa*, *R. sativus* and x*Brassicoraphanus*.

Supplemental Data Set S3. Gene ontology analysis of homoeologous gene pairs showing biased, convergent and constant expression.

Supplemental Data Set S4. Number of read pairs mapped on *B. rapa*, *R. sativus*, and x*Brassicoraphanus* genomes.

## Acknowledgments

Mary Gehring at Whitehead Institute, MIT commented on the manuscript. This work is dedicated to late Dr. Woo Jang-Choon (1898–1959), also known as Nagaharu U in Japanese, for his 60th memorial anniversary.

## Funding

This work was supported by the Next-Generation BioGreen 21 Program (PJ013262) and the National Agricultural Genome Program (PJ013440) by Rural Development Administration (RDA), Republic of Korea.

## Author contributions

J.H.H. conceived the project. H.S., J.E.P., H.R.P. and J.H.H. designed the study. H.S., J.E.P., W.L.C., H.Y.S. and G.Y. performed molecular biology experiments and analyzed the data. J.E.P. performed FACS analysis. H.R.P. and W.K. performed immunolocalization experiments. H.S., S.H.Y., S.K., J.H.A. and J.-S.K. performed bioinformatics analysis. N.E.W., H.R.B. and S.Perumal performed FISH analysis. Y.-M.K. and N.K. performed annotation analysis. K.K. and T.-J.Y. analyzed chloroplast genomes. S.Park, J.A.K., Y.P.L. and S.-S.L. provided plant materials. H.S., J.E.P., H.R.P., W.L.C., S.H.Y., W.K., H.Y.S., J.Y.L., G.Y., T.K., J.K., H.J., D.H.K., Y.S.K., H.-M.J., J.Y. and S.S. prepared plant materials. W.L.C., S.W.Y., J.Y.L., B.-S.P., T.-F.H., T.-J.Y., D.C., H.H.K. and S.-S.L. commented on the manuscript. H.S., J.E.P., H.R.P. and J.H.H. wrote the manuscript with help from all co-authors.

## Conflict of interest statement

None declared.

